# Ecological Factors Affecting Prosociality in Marmoset Monkeys

**DOI:** 10.1101/2025.06.26.661771

**Authors:** Anand R. Mysorekar, Anushka Vispute, Cory Miller

**Affiliations:** Cortical Systems and Behavior Laboratory, University of California, San Diego

## Abstract

Prosocial behaviors, such as cooperation and food sharing, are critical for maintaining group cohesion in social species, yet the influence of transient physiological states on these behaviors remains poorly understood. This study investigates how short-term ecological factors impact social behavior in common marmosets (*Callithrix jacchus*), a highly prosocial nonhuman primate species. Specifically, we tested how food divisibility (small vs. large food items) and time since food access (TSFA; 0 hours, 1 hour, and 3 hours) influenced food sharing behavior. Results revealed that larger food portions consistently promoted prosocial interactions, while longer durations since food access shifted behavior toward individualism. A significant interaction between TSFA and food size indicates that the effect of food size on prosocial behavior varies with hunger state: increasing TSFA weakens the prosocial impact of large food portions. These findings suggest that marmoset prosociality is sensitive to immediate ecological conditions, reflecting flexible, context-dependent social decision-making.

## Introduction

Understanding the conditions that shape prosocial behavior is central to unraveling the evolution of cooperation (de Waal, 2008; Jaeggi & Gurven, 2013). Among nonhuman primates, the prosocial tendencies of common marmosets (*Callithrix jacchus*) have received particular attention because of their high levels of unsolicited food sharing, cooperative care, and social tolerance: traits thought to be related to their cooperative breeding (Burkart et al., 2007; Burkart & van Schaik, 2020). While long-term social structures such as kinship and pair bonds have traditionally been seen as drivers of prosociality (Silk et al., 2013), emerging evidence suggests that transient ecological factors may also exert powerful, immediate effects on cooperative decisions (Yamada, 2017). Yet, the influence of such short-term states, particularly hunger, on prosocial behaviors remains poorly understood (Mustoe et al., 2015; Yamada, 2017).

Although prior research has demonstrated that marmosets routinely share food with others, including unrelated and unfamiliar individuals (Burkart et al., 2007; Mustoe et al., 2015), few studies have experimentally manipulated both resource availability and divisibility to examine their joint impact on cooperative behavior (Brown et al., 2005). Here, we sought to bridge this gap by investigating how short-term ecological variables, specifically, food size and TSFA, modulate food sharing behavior in marmosets.

## Methods

### Subjects

This study involved 29 common marmosets (*Callithrix jacchus*): 12 males (4 parents, 8 offspring) and 17 females (6 parents, 11 offspring) housed across six different family groups. All animals were kept in enriched enclosures on a 12-hour light/dark cycle, with ad libitum access to water and a controlled diet provided outside of experimental sessions. All experimental procedures were approved by the Institutional Animal Care and Use Committee at the University of California, San Diego.

Data collection was conducted in two phases. During the first phase, animals in three cages were tested under the experimental paradigms outlined below. After completing testing in these cages, a method for individual identification was introduced to enable analysis of individual, age, and sex differences. In the second phase, three additional cages were tested using this identification method. Subjects’ hair tufts were temporarily dyed using Arctic Fox Semi-Permanent Hair Color (selected for its vegan, cruelty-free formulation, and non-toxic nature) in unique combinations to allow for identification in video recordings.

### Experimental Design and Procedure

We used a within-cage design to investigate the effects of food size and TSFA on prosocial behavior. Each cage was exposed to three TSFA conditions (0 hours, 1 hour, and 3 hours) and presented with two food sizes (small and large). Each condition was presented 15 times per cage, totaling 90 trials per cage and 540 trials across all six cages. Trial order was randomized and counterbalanced across cages to control for order effects.

Each cage completed 18 test sessions, with six sessions conducted under each TSFA condition. Each session consisted of five trials. At the start of each trial, the food item was placed in the test box with randomized placement to prevent operator bias. The door separating the cage from the entrance tunnel was then opened, allowing monkeys to move freely between the test box and their home cage. A five-minute trial began immediately after food placement and continued until one of the following occurred: the five-minute period ended, the food was fully consumed, or a monkey removed the food and brought it into the home cage. At the end of each trial, the door was closed, and a five-minute inter-trial interval allowed subjects to consume the food and reduced potential carryover effects such as lingering satiety. Sessions were conducted at approximately the same time each day to minimize variability due to circadian rhythms.

The test box measured 61 cm × 47.75 cm × 47.75 cm and was constructed from plexiglass and 80/20 T-Slotted extrusions, allowing for unrestricted visual and physical interaction between subjects. An overhead camera provided a full view of the test box floor for later behavioral coding. Fig. 1a illustrates this setup.

**Fig. 1:**
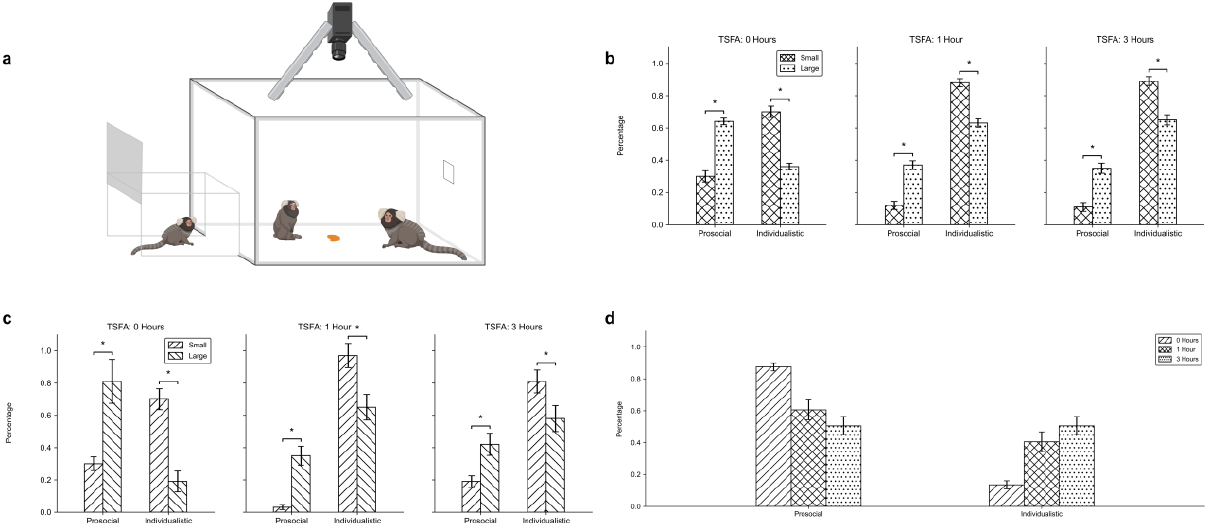
Effects of food size and time since food access (TSFA) on marmoset prosocial behavior a Experimental setup: trials occurred in a test box connected to the home cage. Monkeys could freely enter the test box through the entrance tunnel once the door was opened. An overhead camera recorded all sessions. Food was placed at a randomized location in each trial b Behavior proportions across food sizes at three TSFA conditions in the full dataset (small food: crosshatch; large food: dotted) c Same comparison as in panel b for a thresholded subset of monkeys that participated in all three TSFA conditions with both food sizes (small food: forward slash; large food: backslash) d Behavior proportions across TSFA conditions from a single representative cage (0 h: forward slash; 1 h: crosshatch; 3 h: dotted) Significance markers are indicated with asterisks (*p* < 0.05)

The food used was Strawberry Bio-Serve Electro-Gel - Electrolyte Replenisher, a thermoreversible gel containing balanced concentrations of electrolytes and glucose, commonly used as dietary enrichment for marmosets. Food size was manipulated by presenting large portions as one-fourth of a full gel unit and small portions as one-sixteenth. These sizes were based on prior lab observations indicating they were sufficient for simultaneous consumption by multiple monkeys, enabling food sharing.

### Behavioral Coding and Measures

Videos were analyzed post-experiment to identify distinct behaviors exhibited during the trials. Because no standardized ethogram existed for short-term food sharing in marmosets, we developed one through direct observation of the sessions. This ethogram (Table 1) was then used to manually code all behaviors. Two trained observers independently coded the data: one for the first three cages and the other for the last three. Inter-observer reliability was high, with an intraclass correlation coefficient (ICC, model 2,1) of 0.95 (95% CI: [0.95, 0.95], *p* < 0.001) indicating excellent agreement.

**Table 1:** Categorization of observed behaviors into prosocial, individualistic, and neutral groups based on food sharing interactions.

| Behavior Category | Behavior Name | Description |
| --- | --- | --- |
| Prosocial | Food Sharing | Two or more monkeys eat simultaneously, without conflict. |
| Prosocial | Tolerant Sharing | One monkey eats while another is nearby, without conflict or competition. |
| Prosocial | Sequential Engagement | Monkeys take turns eating; one stops to allow another to eat before resuming. |
| Prosocial | Allofeeding | One monkey directly transfers food to another, either mouth-to-mouth or by offering a portion. |
| Individualistic | Solitary Consumption | One monkey eats alone, with no interaction from others nearby. |
| Individualistic | Snatching | One monkey takes the food and immediately retreats into the home cage. |
| Individualistic | Contest | Two or more monkeys engage in aggressive behavior over the food (e.g., fighting). |
| Neutral | Non-Participation | Monkeys do not interact with the food or engage in any food-related behavior during the trial. |

### Data Analysis

Data were processed using Microsoft Excel and Python, and statistical analyses were conducted with the pingouin package. Repeated-measures ANOVAs were used to assess the main and interaction effects of food size and TSFA on behavioral proportions. Type II sums of squares were applied to emphasize main effects, consistent with our primary hypotheses. To control for cage-level variation in TSFA, secondary analyses were performed using three thresholded datasets with increasing inclusion strictness. All statistical tests were two-tailed with α = 0.05.

## Results

A repeated-measures ANOVA on the full dataset (Fig. 1b) revealed a significant effect of food size on prosocial behavior (*F*(1,5) = 9.66, *p* = 0.027). Across all TSFA conditions, monkeys exhibited more prosocial behavior with large food portions and more individualistic behavior with small portions. Although the effect of food size was present at all TSFA levels, a significant interaction effect (*p* = 0.049) indicates that the strength of this effect varied with hunger state.

To ensure this effect was not driven by sampling bias, we conducted secondary analyses using thresholded datasets with increasingly strict inclusion criteria. The most restrictive dataset (Fig. 1c) included only monkeys that participated in all three time points and both food sizes. In all cases, repeated-measures ANOVAs confirmed a significant food size effect (*p*s = 0.048, 0.046, and 0.049 for the least to most restrictive thresholds, respectively), indicating that this effect was robust to variation in individual participation. TSFA effects were not significant in these restricted datasets.

Separately, a repeated-measures ANOVA on the full dataset revealed a significant effect of TSFA (*F*(2,10) = 8.30, *p* = 0.008), indicating that overall behavior varied across time conditions. However, pairwise comparisons showed that only the contrast between 0 and 3 hours was significant (*p* = 0.047), while comparisons between 0 vs. 1 hour (*p* = 0.054) and 1 vs. 3 hours (*p* = 0.87) were not. This pattern suggests a gradual behavioral shift rather than a stepwise change. Fig. 1d shows a representative cage in which prosocial behavior declined and individualistic behavior increased with greater TSFA. While this pattern was observed in some cages, others remained stable across conditions, likely contributing to the lack of consistent pairwise effects.

## Discussion

We found that marmosets were more likely to share food when portions were large, indicating that resource abundance promotes prosocial behavior. However, during longer TSFA durations, individuals became more self-serving, suggesting that acute hunger reduces cooperation in socially competitive contexts. This aligns with prior findings that resource abundance promotes prosociality. Wittig et al. (2014), for example, found that chimpanzees were more likely to share food when resources were plentiful. Our findings similarly suggest that marmoset food sharing is context-dependent, increasing with larger portions rather than being automatic or unconditional. A key distinction is that while previous studies often emphasize long-term social structures, such as kinship or reciprocity, our results highlight the role of short-term ecological variables, specifically food size and time since food access, as immediate drivers of cooperative behavior.

The findings presented here contrast with work suggesting that hunger may enhance decision-making efficiency in marmosets. Carvalho et al. (2022) found that food-deprived marmosets made more consistent economic choices in a non-social foraging task, maximizing outcomes rather than acting impulsively. In contrast, our results show that acute hunger (longer TSFA) decreased prosocial food sharing and promoted individualism. The divergence likely reflects task context: our study involved competitive food sharing with conspecifics, whereas Carvalho and colleagues (2022) focused on isolated foraging. The cognitive demands and social dynamics differ substantially between these settings, suggesting that the behavioral effects of hunger are not universal, but highly dependent on context.

Our results underscore the importance of both external resource conditions and internal physiological states in shaping cooperative behavior. Larger food portions increased prosocial behavior, while acute hunger appeared to diminish it. This interaction suggests that social decision-making is flexible and influenced by immediate ecological constraints. Neurobiologically, these results motivate future studies investigating how hunger modulates neural circuits involved in reward, impulse control, and social cognition, such as the orbitofrontal cortex and anterior cingulate cortex (Apps et al., 2016).

This study provides novel evidence that prosocial behavior in marmosets is not fixed, but dynamically shaped by short-term ecological and physiological variables. By jointly manipulating food portion size and time since last food access, we demonstrate that cooperation is promoted by resource abundance but suppressed by acute hunger, highlighting the context sensitivity of social behavior even in a highly cooperative species. More broadly, these findings lay the groundwork for future investigations into the neural and hormonal mechanisms that mediate the interplay between internal states and social behavior (Lefevre et al. 2024), advancing our understanding of the neurobiological foundations of cooperation.

## Acknowledgements

This work was supported by grants to CTM from the UCSD Sanford Institute for Compassion and Empathy and the NIH (R01 DC012087). ARM was supported in part by the SURFiN (Shenoy Undergraduate Research Fellowship in Neuroscience) at University of California, San Diego (UCSD). We thank the UCSD animal care staff and research facilities for their support throughout the project.

## Data Availability

Code and aggregated analysis files are available from the corresponding author upon request.

## Notes

### Competing Interest Statement

The authors have declared no competing interest.

